# Low-order assemblies drive oncogenic RTK fusion signaling without condensation

**DOI:** 10.64898/2025.12.10.693466

**Authors:** David Gonzalez-Martinez, Thomas R. Mumford, Delaney Wilde, Sofia Wissert, Yuzhi (Carol) Gao, Emily Brackhahn, Richard Kriwacki, Elizabeth Rhoades, Lukasz J. Bugaj

**Author notes:** equal contributions (David Gonzalez-Martinez, Thomas R. Mumford).

## Abstract

Receptor tyrosine kinase (RTK) fusions are a large class of oncoproteins found in ∼5% of cancers. Key questions remain, however, about how RTK fusions transmit oncogenic signals, including how these largely cytoplasmic proteins activate downstream pathways that originate at the plasma membrane. Fusions are multimeric and can form mesoscale condensates in cancer cells, and condensation has been implicated as an essential mechanism to enable signal transmission from the cytoplasm. However, whether condensates play a causal role, or whether smaller ‘diffuse’ assemblies are sufficient to transduce signals, has been challenging to establish. Here we apply advanced microscopy, single-cell analysis, and synthetic fusions to determine the principles by which multimerization and condensation drive signaling from cytoplasmic RTK fusions. For EML4-ALK, a prominent fusion that forms condensates, we found poor correlation between condensation and signaling. By contrast, EML4-ALK activity was abundant in the diffuse phase, and the kinetics of diffuse-phase activity aligned more closely with downstream Erk signaling than did kinetics of signaling within condensates. Synthetic RTK fusions showed that cytoplasmic ALK or RET fusion dimers—and even constitutively active monomers—were sufficient to induce strong Ras-Erk signaling despite the absence of condensates, and diffuse fusions were sufficient to transform cells in vitro and in subcutaneous tumor models. A panel of various other cancer-driving RTK fusions showed that low-order multimerization was universal across fusions, whereas mesoscale condensation was rare and did not correlate with signaling. Our results suggest that low-order fusion multimerization is sufficient to drive its phosphorylation, which is necessary and sufficient to trigger downstream oncogenic signaling.

## Introduction

RTK fusions are a large class of oncogenes found in ∼5% of cancers across tissue types^1^. Although over 1000 distinct RTK fusions have been identified, these varied proteins share a common composition: the intracellular fragment of a normally transmembrane RTK is fused via chromosomal rearrangements to a partner fragment from a distinct protein^1,2^. Partner fragments are typically homo-oligomeric^3,4^, and oligomerization results in forced proximity between the fusion RTK domains that triggers their phosphorylation and downstream signaling, mimicking the effects of ligands on transmembrane RTKs^2,5–7^. However, unlike transmembrane RTKs, fusions often lack transmembrane domains and therefore localize in the cytoplasm, away from the plasma membrane and the downstream enzymes that reside therein. Nevertheless, fusions retain the ability to drive tonic downstream signaling, although the mechanisms remain unclear. Understanding these underlying mechanisms is critically important to inform therapeutic strategies against this broad range of oncoproteins.

RTK fusions can form mesoscale protein condensates, defined as membrane-less compartments that form microscopically visible assemblies on the 0.1-1 µm length scale^6–10^. This was first shown for EML4-ALK, an RTK fusion that drives ∼5% of non-small cell lung cancer^9,11^. Condensation resulted from two common properties of fusions: 1) homo-oligomerization of the partner fragment (EML4), and 2) activity of the kinase fragment (ALK), which recruits adapters that themselves form multivalent interactions^6^. Notably, other RTK fusions form condensates independent of kinase signaling^6,7^ (**Figure 1A**). Condensation has been proposed as an essential signaling mechanism because condensates could provide a cytoplasmic platform to assemble and concentrate essential signaling factors that would normally assemble at the membrane. Indeed, several studies have now shown that signaling adapters and enzymes are recruited to fusion condensates^6,7,10,12^. On the other hand, a recent screen of oncogenic fusions found that only a small percentage of oncogenic RTK fusions robustly form condensates, raising questions about the necessity of condensation for fusion signaling^4^. Overall, the causality of condensation in downstream signaling has been challenging to determine because low-order fusion multimerization drives both condensation and kinase activity, and, in the case of EML4-ALK, kinase activity further promotes condensation. Therefore, perturbations of condensation are frequently also perturbations of kinase activity, confounding conclusions on the nature of the essential signaling unit and on the specific role of condensates^6^.

**Figure 1.**
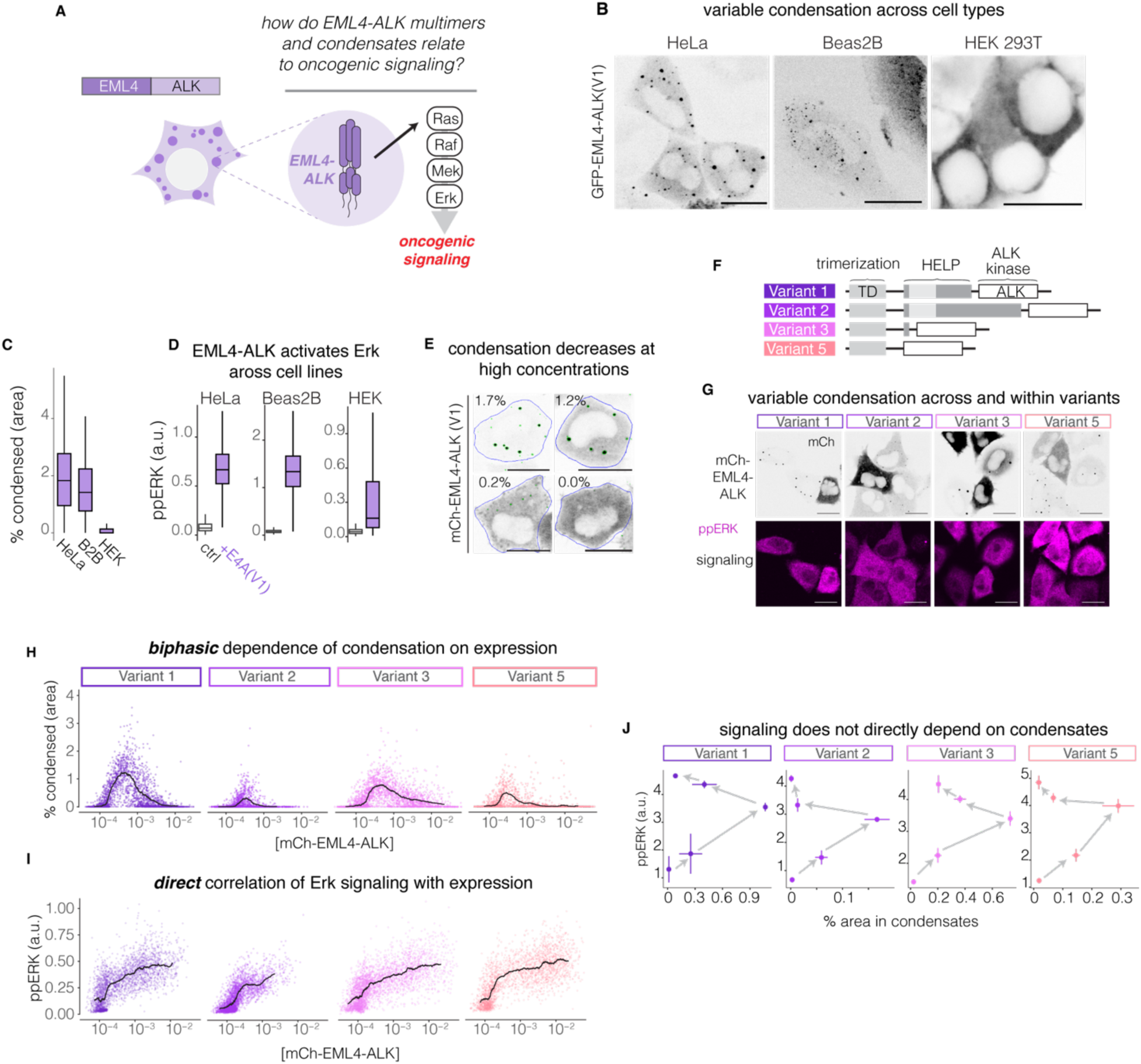
EML4-ALK signaling does not correlate with condensation A) Dissecting the role of condensation in EML4-ALK signaling. B) Representative images of HeLa, Beas2B, and HEK 293T cells expressing exogenous GFP-EML4-ALK(V1). Scale bars = 20 µm. C) Percent cell area in EML4-ALK (V1) condensates. n = 838 (HeLa), 355 (Beas2B), 1066(HEK 293T) cells. D) Erk phosphorylation in HeLa, Beas2B, and HEK 293T cells with (purple) or without (gray) expression of exogenous mCh-EML4-ALK(V1). n = 3427, 1124 (HeLa); 2197, 555 (Beas2B); and 13454, 5067 (HEK 293T) for control and EML4-ALK(V1)+, respectively. E) Representative images of condensate segmentation and condensate heterogeneity in HeLa cells expressing mCh-EML4-ALK (V1). Numbers indicate percent of cell area (pixels) classified as condensates. Scale bars = 20 µm. F) Domain representation of EML4-ALK V1, V2, V3, and V5. G) Representative images of mCh-EML4-ALK and respective pERK immunostaining in HeLa cells expressing the indicated variants. Scale bars = 20 µm. H) Percent of cell area containing condensates in HeLa cells expressing mCh-EML4-ALK variants as a function of mCh intensity. Data points represent single cells. Solid line represents rolling average. I) pERK intensity in HeLa cells expressing mCh-EML4-ALK variants as a function of mCh intensity. Data points represent single cells. Solid line represents rolling average. For (H,I) n = 2331 (V1), 2844 (V2), 3088(V3), 2300(V5). J) pERK intensity as a function of mCh-EML4-ALK condensation. Arrows point towards bins of increasing levels of expression. Data points represent mean +/-SEM of three replicates per bin. For (J) n = 343-1065 (V1); 51 - 1678 (V2); 399 - 1160 (V3); 243 - 754 (V5) cells per for each bin.

Here we overcome these challenges to determine how RTK fusion multimerization and condensation relate to signaling and oncogenic potential of RTK fusions (**Figure 1A**). Using quantitative live-and fixed-cell microscopy, single-cell and single-condensate analysis, and a panel of natural and synthetic fusions, we find that phosphorylation downstream of kinase multimerization is necessary and sufficient to trigger signaling downstream of RTK fusions, whereas condensation is dispensable. Our work establishes general principles of RTK fusion signaling and highlights the importance of low-order, submicroscopic fusion assemblies in oncogenic signal transduction.

## Results

### EML4-ALK signaling does not correlate with condensation

To understand how condensation relates to downstream signaling, we first sought a suitable cell model. Expression of EML4-ALK variant 1 (V1) in multiple cell types revealed substantial heterogeneity in condensate formation. Condensation — quantified as the percentage of pixels classified within condensates^13^ — was strongest in HeLa cells, somewhat lower in lung epithelial Beas2B cells, and weakest in HEK 293T cells (**Figure 1B,C**). We therefore used HeLa cells for the majority of experiments in this study. Notably, despite differences in condensation, EML4-ALK drove strong ERK activation across all cell lines tested (**Figure 1D, S1**).

We next examined the relationship of EML4-ALK to downstream signaling on a single-cell level. Condensation was heterogeneous between individual cells, with a notable decrease in condensation in cells with higher expression (**Figure 1E**). We recognized that this heterogeneity could be leveraged to test the hypothesis that condensates are causal for signaling, since cells with more condensates should show a concomitant increase in signaling. We examined this relationship across four common variants of EML4-ALK (V1, 2, 3, and 5) found in patient cancers, all of which have the capacity to condense,^6,10,12^ albeit to different extents (**Figure 1F,G**)^14–16^. In HeLa cells, V1 and V3 formed condensates in most cells and on average formed the largest condensates, whereas V2 and V5 condensed in fewer cells, consistent with prior results (**Figure 1G**)^6,10,12^. However, across all variants, condensation was heterogeneous between cells and was most prominent in low/intermediate expressers, whereas high expressers appeared diffuse. Notably, ERK activity was not restricted to cells with condensates and on the contrary appeared stronger in cells with strong, diffuse expression of EML4-ALK (**Figure 1G**). For all variants, condensation showed a biphasic relationship with EML4-ALK expression, where condensation was maximal at low-intermediate levels of oncogene and decreased beyond this point (**Figure 1H**). In stark contrast, Erk signaling correlated directly and monotonically with EML4-ALK expression (**Figure 1I**). Therefore, there was no direct relationship between condensates and Ras-Erk signaling (**Figure 1J**); the cells that exhibited the highest Erk signaling levels most frequently showed only diffuse expression of EML4-ALK.

### Amplification of EML4-ALK condensation does not increase ERK signaling

To further test for a causal role of condensates on EML4-ALK signaling, we sought methods to increase condensate size and assess the effect on Erk signaling. The biphasic dependence of condensation on concentration suggested a limiting component within the heterotypic EML4-ALK condensates^17,18^. We thus asked whether condensation could be increased by supplementation with GRB2, a multivalent adaptor whose interaction is essential for EML4-ALK condensation (**Figure 2A**)^6^. Indeed, co-expression of GRB2-GFP augmented EML4-ALK condensation across expression levels, including at high expression where condensation was otherwise absent. Increased condensation was observed for all variants, although the magnitude and concentration-dependence was unique to each variant (**Figure 2B-D**).

**Figure 2.**
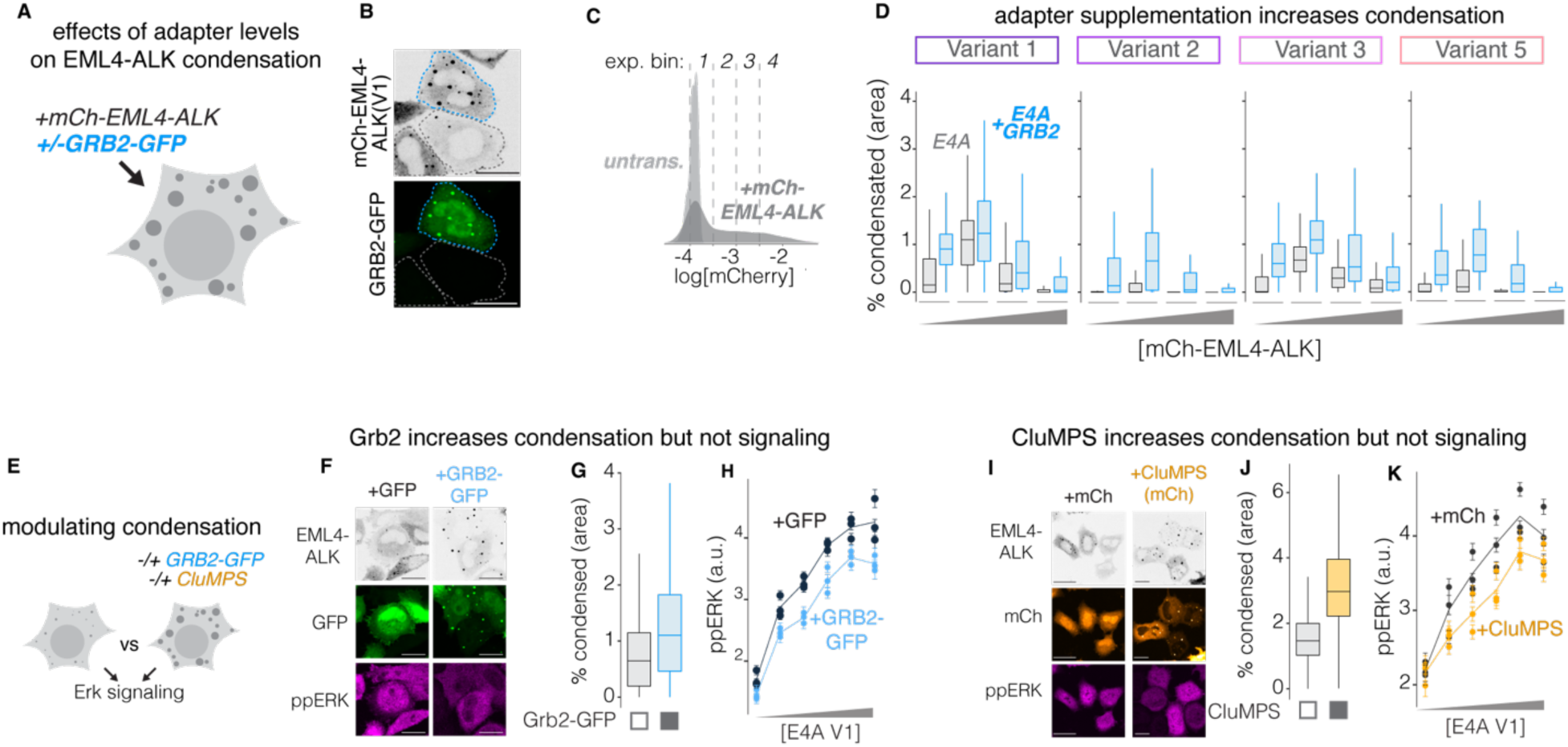
Amplifying EML4-ALK condensation does not increase signaling. A) Investigating the effect of Grb2 supplementation on EML4-ALK condensation. B) Representative images of HeLa cells expressing mCh-EML4-ALK(V1) with (blue outline) or without (gray outline) GRB2-GFP co-expression. Scale bars = 20 µm. C) Representative expression distribution and binning strategy for mCh-EML4-ALK expressed in HeLa cells. The same absolute fluorescence thresholds were applied to all variants. D)Percent cell area with condensates in HeLa cells co-expressing mCh-EML4-ALK variants with GRB2-GFP (blue) or mCh-EML4-ALK(V1) alone (gray), as a function of the mCh binned expression. n = 388-717, 67-398 (V1); 51-1678, 12-573 (V2); 399-1160, 80-292 (V3); 245-754, 41-251(V5) cells for E4A alone and GRB2-GFP co-expression respectively. Boxplots show median and upper/lower quartile, and whiskers extend to 1.5*IQR. E) Investigating effects of condensate modulation through co-expression of either GRB2-GFP or CluMPS. F) Representative images of cells from (G,H). Scale bars = 20 µm. G) Degree of condensation of mCh-EML4-ALK(V1) in HeLa cells co-expressing GRB2-GFP (blue) or GFP alone (grey). n = 2333 (+GFP only) and 2215 (+GRB2-GFP). H) pERK intensity of HeLa cells co-expressing mCh-EML4-ALK(V1) with GRB2-GFP (blue) or GFP alone (black) as a function of mCh-EML4-ALK binned expression. Datapoints represent mean +/-SEM of 67-198 (+GFP) and 32-262 (+GRB2-GFP) cells. I) Representative images of cells from (J,K). Scale bars = 20 µm. J) Degree of condensation of GFP-EML4-ALK(V1) in HeLa cells co-expressing a CluMPS probe (LaG17-mCh-HOTag3, orange) or mCh alone (grey). n = 3606 (+mCh) and 3431 (+CluMPS) cells. K) pERK intensity of HeLa cells co-expressing GFP-EML4-ALK(V1) with mCh (black) or CluMPS (yellow), as a function of GFP-EML4-ALK(V1) binned expression. Data points represent the mean +/-SEM of 71-291 (+mCh) cells and 122-302 (+CluMPS) cells. H,K) Line represents the mean of 3 replicates. H,K) Cells were gated into bins using quarter log steps in fluorescence.

While condensation was elevated in GRB2-GFP-expressing cells, ERK signaling was not increased at any level of mCh-EML4-ALK (V1) expression (**Figures 2F-H**). These data indicate that condensation does not enhance signaling. However, a potential confounding factor is that GRB2 overexpression could alter the stoichiometric balance of proteins required for efficient signaling from fusion assemblies. Therefore, we tested an independent method to enhance clustering. We co-expressed GFP-EML4-ALK with a CluMPS reporter that targets GFP^13^ (**Figure 2E**). CluMPS is multimeric and, upon binding a multimerized target, enlarges the size of target assemblies. As with Grb2, increased condensation due to CluMPS co-expression did not increase signaling at any level of GFP-EML4-ALK concentration (**Figure 2I-K**). Together, these data do not support condensation as a driver or potentiator of downstream signaling.

### Diffuse-phase EML4-ALK activity corresponds to downstream signaling

The lack of correspondence between condensation and Ras-Erk signaling suggested that signaling may primarily emanate from EML4-ALK in the diffuse phase. To investigate this possibility, we measured ALK phosphorylation (pALK) in HeLa cells that expressed EML4-ALK(V1) (**Figure 3A)**. Immunostaining for pALK revealed strong correlation between total pALK and EML4-ALK expression, but poor correlation between total pALK and condensation (**Figure 3 B-D**), paralleling our results for Erk signaling. We next separately analyzed the pALK signal corresponding to the condensed vs diffuse compartments (**Figure 3E**). Although the mean pixel intensity of pALK was higher in the condensed phase and increased with EML4-ALK expression level (**Figure 3F**), the total integrated pALK signal was higher in the diffuse phase due to its larger volume fraction (**Figure 3G,H**). In addition, at the highest expression levels where condensation decreased, the integrated pALK signal in condensates plateaued and decreased as a fraction of total pALK. By contrast, the diffuse integrated pALK signals continued to rise, in concert with pERK (**Figure 3G,H**, **Figure 1I**).

**Figure 3.**
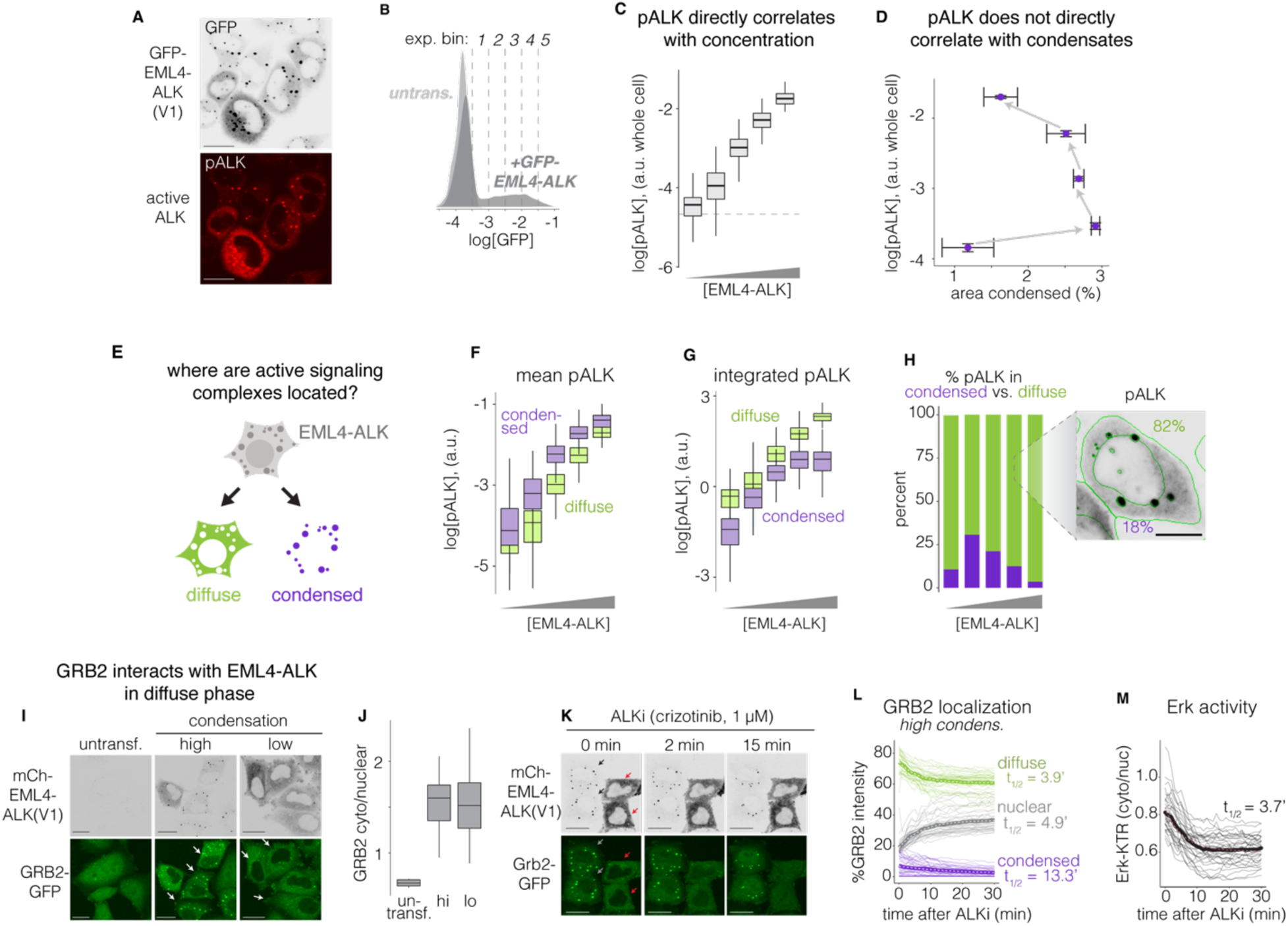
Dilute-phase activity of EML4-ALK(V1) corresponds to downstream Erk signaling. A) Representative images GFP-EML4-ALK(V1) and pALK immunostaining in HeLa cells. Scale bars = 20 µm. B) Expression distribution and binning strategy for mCh-EML4-ALK expressed in HeLa cells. C) Whole-cell pALK intensity of HeLa cells expressing GFP-EML4-ALK(V1) as a function of expression level. Horizontal line represents background pALK signal from untransfected HeLa cells. D) pALK intensity as a function of GFP-EML4-ALK condensation. Arrows point towards bins of increasing levels of expression. E) Separately analyzing pALK in the diffuse vs condensed phase. (F) Mean and (G) integrated pALK intensity of pixels corresponding to condensed (purple) or diffuse (green) GFP-EML4-ALK in HeLa cells, as a function of GFP-EML4-ALK expression. H) Fraction of pALK intensity in pixels corresponding to condensed and diffuse GFP-EML4-ALK(V1) as a function of expression. Inset: representative image of a cell from fusion expression bin 4. Scale bars = 10 µm. For (C-H) n = 53-299 cells per bin. I) Representative images of mCh-EML4-ALK and GRB2:GFP (endogenous) in HeLa cells, highlighting representative cells with high condensation (> 0.2% area) or low condensation (< 0.2% area). Scale bars = 20 µm. White arrows show nuclear exclusion of GRB2 in cells expressing EML4-ALK, indicating interaction between GRB2 and EML4-ALK in the cytoplasm. J) Quantification of the ratio of cytoplasmic to nuclear fluorescence of GRB2:GFP. N = 76 (untransfected), 45 (low), 81 (high). K) Representative images of GRB2:GFP localization in cells expressing mCh-EML4-ALK(V1) subsequent to treatment with ALKi (crizotinib, 1µM). Scale bars = 20 µm. Black and red arrows show nuclear exclusion in high-and low-condensation cells, respectively. L) Percentage of total GRB2:GFP fluorescence found in the nucleus (gray), condensates (purple), or diffuse in the cytoplasm following ALKi treatment in high condensation cells. Low condensation cells are quantified in **Figure S2A**. M) Quantification of Erk activity using the ERK-KTR reporter in the same cells as in (L). Data from low condensation cells is quantified in **Figure S2B**. In (L,M), transparent traces represent single cells. Solid trace represents population mean. Dashed traces represent fit of the mean to a single exponential decay, which was used to calculate t_1/2_.

As a further test, we examined the timescales of changes of ALK activity in the different compartments by measuring adapter localization and ERK signaling after kinase inhibition. EML4-ALK activity leads to its phosphorylation and binding by GRB2. We thus reasoned that GRB2 could serve as a biosensor that reports on the location of ALK activity. We measured the localization of endogenously-tagged GRB2 in HeLa cells (HeLa GRB2:GFP) and compared its response to that of ERK signaling (Erk-KTR)^19^. In the absence of EML4-ALK, GRB2 was found in both the cytoplasm and the nucleus. Upon expression, EML4-ALK (V1) bound GRB2 and sequestered it in the cytoplasm, in both the diffuse and condensed compartments (**Figure 3I**). The cytoplasmic/nuclear ratio of GRB2 was comparable in cells with either low (< 0.2% condensed area) or high (>0.2% condensed area) EML4-ALK condensation (**Figure 3J**). We then treated cells with an ALK inhibitor (ALKi: crizotinib, 1 µM) (**Figure 3K, Supplementary Movie 1)**. In cells with condensates, treatment resulted in 1) dissociation of GRB2 from condensates, 2) decrease of diffuse cytoplasmic GRB2, 3) increase in nuclear GRB2, and 4) decrease in ERK activity. GRB2 dissociated from condensates with a half-life of ∼13 min, in close agreement with a prior report in Beas2B cells^12^. By contrast, diffuse cytoplasmic GRB2 decreased with substantially faster kinetics (t_1/2_ = 3.9 min) (**Figure 3K,L, S2**). If condensates were a key determinant of downstream signaling, the loss of ERK activity should follow the slower kinetics of condensate-associated GRB2. Instead, ERK signaling decreased rapidly (t_1/2_ = 3.7 min), inconsistent with a dependence on condensates but consistent with a dependence on the fast-decaying diffuse cytoplasmic compartment (**Figure 3M**). In low-condensation (mostly high-expressing) cells, loss of diffuse GRB2 also occurs on a comparable timescale as loss of ERK signaling, although both occur slightly slower relative to high-condensation cells, likely due to the higher concentration of EML4-ALK in these cells (**Figure S2**).

### Synthetic ALK fusions show that diffuse-phase, low-order assembly is sufficient for Ras-Erk signaling

To better understand the principles of multimerization, condensation, and signaling, we constructed a panel of synthetic RTK fusions whose multimerization we could explicitly define. For EML4-ALK, The EML4 fragment provides multimerization that is required for both ALK activity and condensation, but the degree of multimerization is neither well-defined nor easy to modulate and is further complicated by potential interactions of EML4 with other species in the cell^20^. We thus investigated the behavior of custom ALK fusions where EML4 was replaced by GFP fused to synthetic coiled-coils that form multimers of defined valency. These allowed us to examine fusions that formed monomers (1mer), dimers (GCN: 2mer), or hexamers (HOTag3: 6mer), and to compare their condensation and signaling to EML4-ALK (**Figure 4A**)^21–23^. To confirm the relative sizes of the signaling species in cells, we performed in-cell fluorescence correlation spectroscopy (FCS), which uses microscopy to measure the timescales of diffusion and the brightness of fluorescent species (**Figure 4B**). We found progressively slower diffusion and increasing brightness as fusion multimerization increased from the 1mer to the 2mer and 6mer, confirming that fusion multimerization in cells was behaving as designed **(Figure 4C-E**).

**Figure 4.**
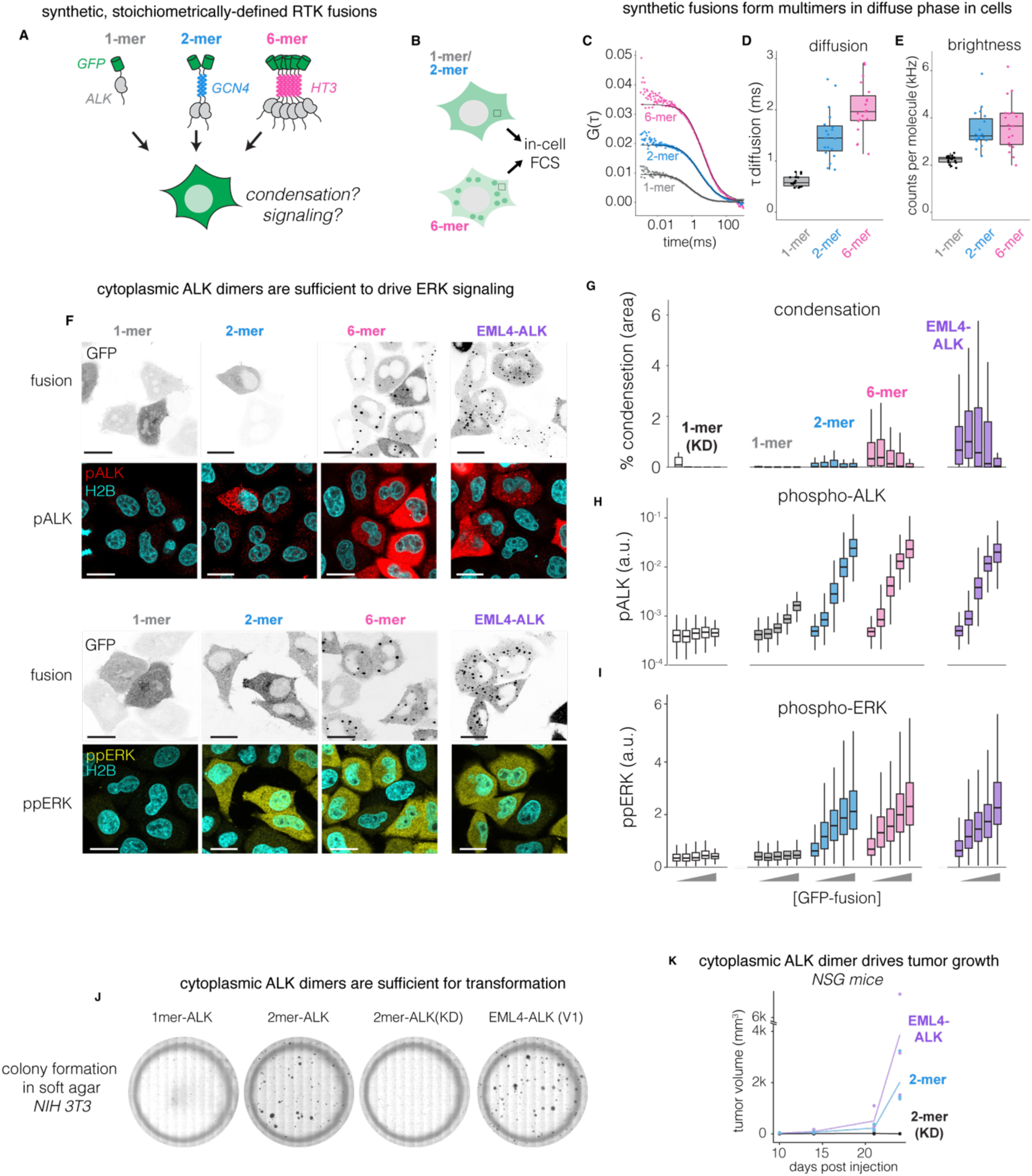
Synthetic RTK fusions show that small, diffuse assemblies are sufficient for ALK signaling. A) Testing synthetic, low-order ALK fusions for propensity to condense and trigger signaling. B) Assessing oligomerization of synthetic ALK constructs using fluorescence correlation spectroscopy (FCS). C) Representative autocorrelation curve of single point measurements in HEK 293T cells expressing 1-mer (gray), 2-mer (blue), or 6-mer (pink). Diffusion times (D) and molecular brightness (E) extracted from fitted autocorrelation curves (black line in E) for 1-mer, 2-mer, or 6-mer ALK fusions expressed in HEK 293T cells. Data represent measurements from 5 cells per construct and are representative of two independent experiments. F) Representative images of pALK or pERK immunofluorescence from HeLa cells expressing 1-mer, 2-mer, 6-mer, or GFP-EML4-ALK(V1). Scale bars = 20 µm. Percent of cell area containing condensates (G), pALK intensity (H), or pERK intensity (I) of the indicated constructs expressed in HeLa cells, as a function of fusion expression. n = ∼50-1000 cells per bin for condensation data and n = ∼200-2000 cells per bin for signaling data.J) Colony formation in soft agar to assess transformation potential of synthetic fusions expressed in NIH 3T3 cells. Images taken after 4 weeks of growth in 0.3% agar and staining with crystal violet. Images are representative of 3 replicates. K) Tumor volume of 3T3 cells stably expressing GFP-EML4-ALK(V1), GFP-2-merALK, or GFP-2-merALK (KD) following subcutaneous injection in NSG mice. Volume was calculated as described in the methods.

In HeLa cells, the synthetic fusions showed a graded increase in condensation as partner valency increased. While condensation was absent for the 1mer and mostly absent for the 2mer, 6mer ALK showed robust condensation, approaching levels observed for EML4-ALK (**Figure 4F,G**). Synthetic fusions demonstrated a biphasic dependence of condensation on concentration, indicating that this biphasic trend is not limited to EML4-ALK and is a general feature of RTK fusions (**Figure 4G**).

Dimerization was sufficient to activate ALK and downstream Erk signaling to levels comparable to 6mer-ALK and EML4-ALK, even in the lowest expression bins, despite its inability to drive condensation (**Figure 4F,H,I**). Monomeric ALK did not transmit signal to ERK, although it could trigger moderate pALK at its highest (likely non-physiological) expression levels. To independently confirm the ability of 2mer-ALK to signal, we expressed 1mer-and 2mer-ALK, as well as a kinase-dead 2mer-ALK (KD, D1270N^24^), in HeLa Grb2:GFP cells and measured the degree of Grb2 sequestration in the cytoplasm (**Figure S3**). Consistent with the Erk signaling results, the 2mer-ALK showed an increased cytoplasmic/nuclear ratio of Grb2 with increasing expression, while the 1mer and the 2mer-ALK(KD) did not.

To confirm our results were not unique to the ALK tail, we constructed an analogous panel of synthetic fusions with the intracellular fragment of RET, the RTK with the second highest prevalence in oncogenic fusions after ALK^1^ (**Figure S4**). RET fusions behaved qualitatively similarly to ALK fusions, showing biphasic condensation and the ability for dimeric RET to signal strongly in the absence of condensation, though minor differences in the magnitudes of condensation and signaling between the ALK and RET fusions were observed.

Signaling from condensate-deficient synthetic fusions was sufficient to transform cells. NIH 3T3 cells that stably expressed 2mer-ALK showed robust colony formation in soft agar, comparable to cells that expressed EML4-ALK, whereas the cells expressing 1mer-ALK and 2mer-ALK(KD) formed no colonies (**Figure 4J**). Additionally, 2mer-ALK-expressing NIH 3T3 cells formed tumors when implanted subcutaneously in immunocompromised mice (**Figure 4K**). Therefore, diffuse-phase fusion dimers are sufficient to trigger oncogenic signaling and transform cells.

### Monomers are sufficient for fusion signaling in the presence of strong phosphorylation

The role of condensation has been challenging to define because fusion multimerization drives both fusion phosphorylation and condensation, and therefore it has been difficult to find perturbations of condensation that do not also interfere with phosphorylation. We therefore sought to generate a synthetic fusion where phosphorylation could be retained in the absence of multimerization and thus in the absence of potential for condensation. We created a constitutively active (CA) ALK monomer (GFP-ALK), hypothesizing that introduction of a kinase-activating point mutation found in oncogenic mutants of transmembrane ALK (CA, F1147L^25^) might permit constitutive phosphorylation of the cytoplasmic fusion (**Figure 5A**). Indeed, 1mer-ALK(CA) showed a strong concentration-dependent increase in ALK phosphorylation, unlike the WT 1-mer (**Figure 5 B-D**). Furthermore, 1mer-ALK(CA) successfully increased downstream pERK to levels approaching those achieved by EML4-ALK, albeit weaker at lower expression levels (**Figure 5C,E**). Notably, neither the WT nor CA 1-mer formed condensates. Signaling of 1mer-ALK(CA) was confirmed by increased C/N ratio of Grb2 in Grb2:GFP HeLa cells (**Figure 5F,G**). FCS confirmed that the WT 1mer, 1mer-ALK(CA), and 1mer-ALK(KD) were indistinguishable in terms of diffusion and brightness, indicating that signaling-competent constructs were not forming cryptic small multimers or condensates within cells (**Figure 5H-J**). In the same manner, constitutively active monomers of the RET intracellular domain (CA, M918T)^26^ could also trigger downstream signaling. 1mer-RET(CA) showed robust kinase activation and was able to activate downstream signaling with comparable magnitude to fusion oncoprotein CCDC6-RET while showing fully diffuse expression (**Figure S5**).

**Figure 5.**
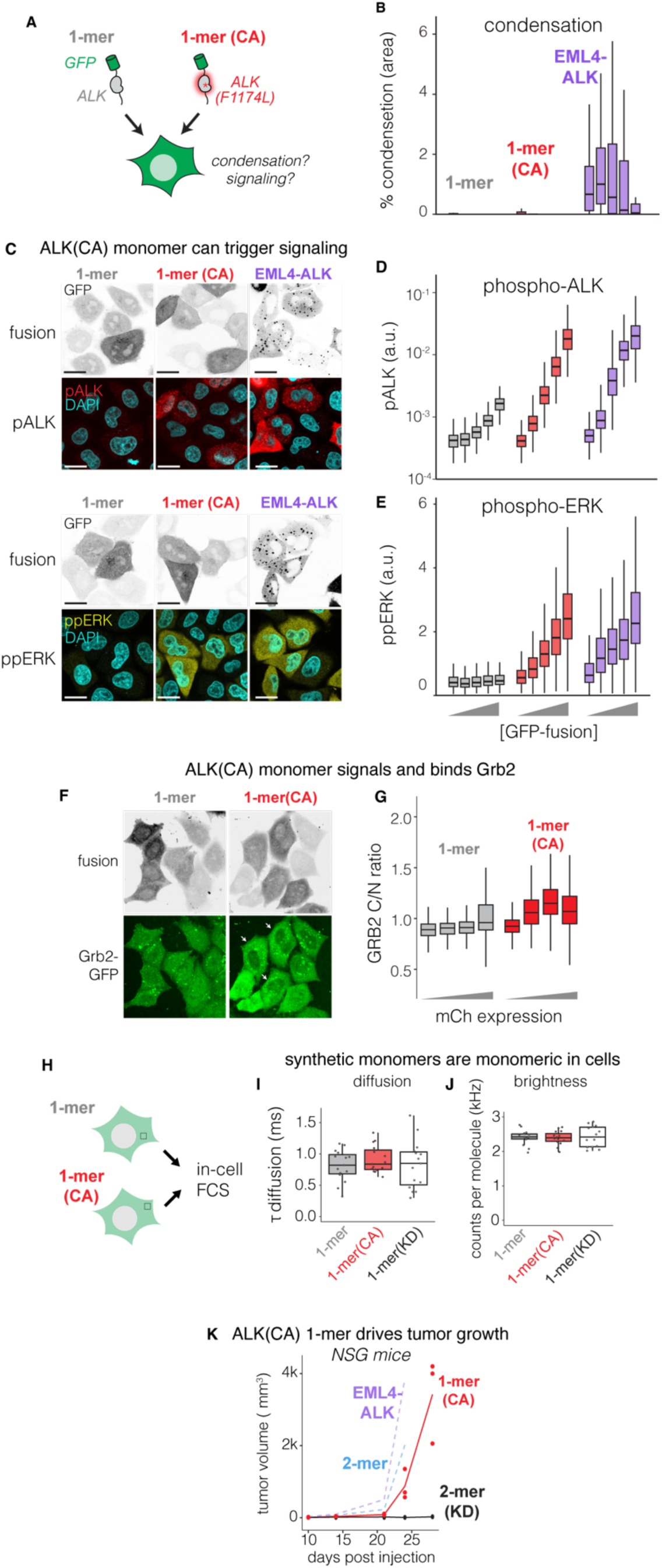
Monomeric but constitutively active ALK kinase domain can activate RAS/ERK signaling. A) Testing the necessity of fusion multimerization with a constitutively active (CA) ALK 1-mer. B) Condensation levels of the 1-mer, 1-mer CA, or GFP-EML4-ALK as a function of expression level. n = ∼50-1000 cells per bin. C) Representative images of the 1-mer, 1-mer (CA), or GFP-EML4-ALK(V1) as well as resultant pALK and pERK levels. Scale bars = 20 µm. Quantification of pALK (D) and pERK (E) levels as a function of expression of the indicated constructs. n = ∼200-2000 cells per bin. F) Representative images of 1-mer and 1-mer(CA) and endogenous GRB2:GFP in HeLa cells. Arrows highlight GRB2:GFP cytoplasmic distribution in cells expressing 1-mer(CA). Scale bars = 20 µm. G) Quantification of GRB2 distribution from (F). n = 398-492 (1-mer), 342-792 (1-mer CA) cells. H) Assessing 1-mer multimerization state through FCS. Diffusion times (I) and molecular brightness (J) extracted from fitted autocorrelation curves of measurements taken from HEK 293T cells expressing the indicated constructs. Data represent measurements from 5 cells per construct and are representative of two independent experiments. K) Tumor volume of 3T3 cells stably expressing GFP-EML4-ALK(V1), GFP-2-merALK, GFP-2-merALK (KD), or 1-merALK (CA) following subcutaneous injection in NSG mice. Volume was calculated as described in the methods. EML4-ALK, 2-mer, and 2-mer(KD) days 10-24 were replicated from Figure 4K.

Finally, we asked if the monomeric, condensate-deficient 1mer-ALK(CA) could drive sufficient signaling to promote tumor growth. Indeed, NIH 3T3 cells expressing this construct formed subcutaneous tumors in immunocompromised mice, albeit with slightly slower kinetics relative to EML4-ALK and 2mer-ALK, consistent with the lower level of signaling from the 1-mer(CA) (**Figure 5K**). The ability of active yet monomeric RTK fusions to drive oncogenic signaling from the cytoplasm identifies phosphorylation of the kinase tail as the critical signaling step required to achieve downstream Ras/Erk signaling. While in cancer-associated fusions, RTK phosphorylation is driven by partner domain multimerization, multimerization itself is not strictly required.

### Low-order multimerization is a widespread feature of oncogenic RTK fusions

To assess whether the principles we describe for EML4-ALK apply broadly across the diverse class of RTK fusions, we analyzed multimerization, condensation, and signaling in a panel of 7 additional cytoplasmic fusions found in cancer, with representation from a variety of common RTK and partner domains (EML4-ALK, PPP1CB-ALK, CCDC6-RET, NCOA4-RET, GSN-NTRK1, TPM3-NTRK1, ETV6-NTRK3, TPM3-ROS1)^4^. Expression of N-terminal GFP fusions of these proteins in HeLa cells revealed a wide variability in condensation. EML4-ALK(V1) and ETV6-NTRK3 showed robust condensation, consistent with previous reports^6,7,9,12^, as did NCOA4-RET (**Figure 6A,B**). However, the remaining fusions appeared mostly diffuse. As observed for EML4-ALK and our synthetic fusions, condensation showed a biphasic relationship to fusion concentration for all fusions (**Figure 6A,B**), whereas Erk signaling showed a direct and monotonic relationship (**Figure 6C**). TPM3-ROS1 was the lone exception to this rule, showing a biphasic relationship between Erk signaling and concentration. Nevertheless, this biphasic relationship was distinct from its biphasic trend for condensation, consistent with a lack of correspondence between condensation and signaling.

**Figure 6.**
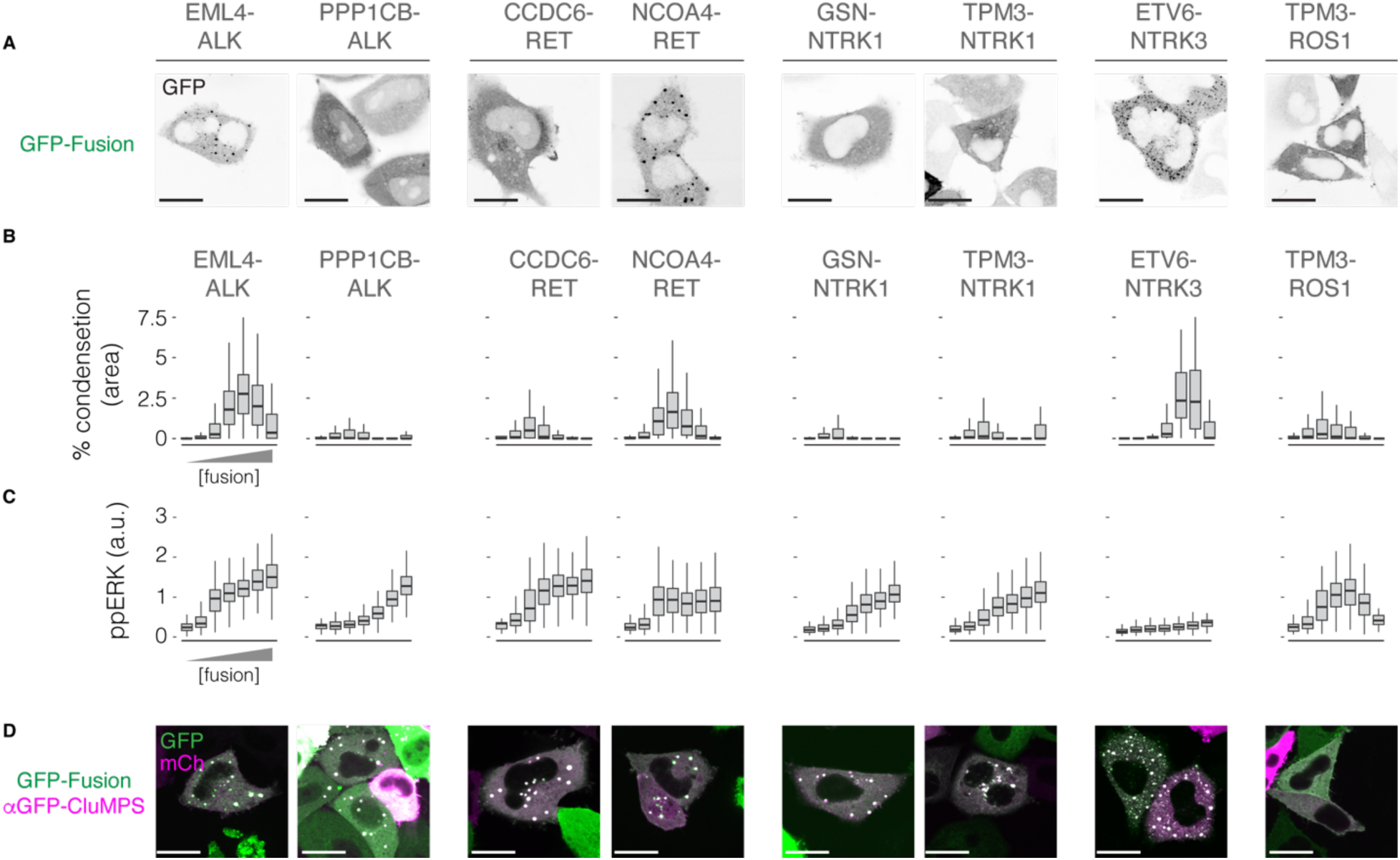
Lack of correlation between condensation and downstream signaling is widespread among oncogenic RTK fusions. A) Representative images of RTK fusions expressed in HeLa cells. Scale bars = 20 µm. Condensation (B) and pERK (C) of RTK fusions in HeLa cells as a function expression. n = ∼50-1200 cells per bin. D) Representative images of HeLa cells co-expressing RTK fusions and CluMPS. Scale bars = 20 µm.

Finally, to determine whether the fusions in our panel formed at least low-order, submicroscopic assemblies, we co-transfected each fusion with a CluMPS reporter that targets GFP. The formation of CluMPS condensates revealed assemblies for all but TPM3-ROS1 (**Figure 6D**). The TPM3 domain has been shown to form dimers which, while sufficient for signaling, can be below the detection limit of CluMPS probes^13,27^. Together, these data indicate that low-order multimers are sufficient to drive oncogenic signaling across the broad class of oncogenic RTK fusions.

## Discussion

Our results challenge the current understanding of the requirements for oncogenic signaling from cytoplasmic RTK fusions. We propose a model where the key determinant for signaling is strong phosphorylation of the RTK fragment. In oncogenic fusions, this is achieved primarily by multimerization of the partner domain, permitting constitutive cross-phosphorylation of neighboring RTK fragments. While multimerization can also trigger formation of mesoscale condensates, we find no evidence that condensates contribute to oncogenic signaling.

Our study leverages single-cell analysis and cell-to-cell heterogeneity to best understand the context-dependence of condensate formation and its impact on signaling, including the roles of fusion concentration, cell type, and levels of downstream adapters like GRB2. The importance of GRB2 for condensation agrees with prior biochemical studies and may also underlie differences in condensation between cell types, though heterogeneity of other confirmed binding adapters (e.g. GAB1, SHC1) likely also contributes^6^. Pixel-level analysis of live-and fixed-cell imaging showed that active fusion signaling complexes exist in the diffuse phase, and that activity in this phase tracks more closely with downstream signaling compared to activity in the condensed phase. The importance of small, diffuse-phase complexes parallels findings from other fields where the function of mesoscale condensation is not clear^28–30^.

While small multimers of the partner fragment were sufficient for RTK fusion signaling, our study finds that partner-mediated multimerization is not necessary, given an alternate route to achieve sufficient phosphorylation of the RTK tail. Nevertheless, partner multimerization is ubiquitous among RTK fusions^4,31–33^. We speculate this is because multimerization is a highly accessible and potent mode of activation: it can be readily obtained across a variety of modular domains during chromosomal rearrangements, and it provides strong, tonic activity of the RTK fragment.

Our conclusion that condensates are dispensable for signaling diverges from previous studies, which suggested a causal role for condensates. One prior study found that EML4-ALK(V5) had diminished capacity to form condensates and linked this effect with decreased Erk signaling^6^. While our study confirms decreased condensation of V5, we found that V5 can signal with ∼equal strength to V1 at equivalent expression levels. This discrepancy may be explained by cell-to-cell variability associated with transient transfection and our ability to correlate expression and signaling at the single cell level. Two other studies found that GFP-tagged CCDC6-RET formed condensates, though only in ∼5-16% of cells^4,34^. Our study similarly finds measurable yet ultimately low levels of condensation for CCDC6-RET, and we show that the remaining majority of non-condensed cells can signal strongly, indicating that condensation is not necessary for CCDC6-RET signaling.

One limitation of our study is that we do not examine fusions within their host cancer cells. Such cells would have undergone selective pressures in the tumor that would yield unique expression profiles that are absent from our studies. Nevertheless, our comprehensive mapping of the expression landscape provides a systematic understanding that could be applied to better understand any number of cancer cells whose oncogenes fall somewhere along the possible expression spectrum.

Our results call for further studies of a fundamental question: how can a cytoplasmic RTK fusion signal through oncogenic pathways that normally originate from the membrane? Our study narrows the set of possible mechanistic explanations that will be the subject of future studies on this important class of oncogenes

## Resource availability

Lead contact

Requests can be made to the lead contact, Lukasz Bugaj

## Materials availability

Plasmids from this manuscript are available upon request from the lead contact.

## Data and code availability

Data generated for this study are available from the lead contact upon request. Code to fit clusters throughout this manuscript is available on Github: https://github.com/BugajLab/Cluster-Fitting

## Supporting information

Supplementary Movie 1. GRB2 localization and Erk activity after ALKi treatment.

## Acknowledgments

This work was supported by the National Institutes of Health (R35 GM138211 for L.J.B. and D.G.M.; P30CA021765, R01 CA246125 to R.K.; R01 NS120625 to E.R.), the American Cancer Society (RSG-22-176-01-TBE to L.J.B.), the National Science Foundation (GRFP to D.W.), St. Jude Research Collaborative Research on the biology and biophysics of RNP granules (R.K), and ALSAC (R.K.).

## Author contributions

D.G.M., T.M., and L.J.B. conceived the study. D.G.M., and T.R.M., E.B., Y.G., D.W., S.W., performed experiments. R.K. provided library of RTK fusions plasmids used in Figure 6. D.G.M., T.M., and E.B. analyzed data. E.R., L.J.B. provided experimental feedback and guidance. L.J.B. supervised the work. D.G.M., T.M. and L.J.B. wrote the manuscript and made figures with editing from all authors.

## Methods

### Cell lines and cell culture

Cell lines were incubated at 37°C and 5% CO_2_ using a standard cell culture incubator. HeLa and HEK 293T (TakaraBio, #632180) cells were cultured using DMEM media (Corning, 10-013-CV) supplemented with 10% fetal bovine serum (FBS) and 1% penicillin/streptomycin (P/S) (Gibco, 15140-122). Beas2B cells were cultured using RPMI-1640 (Corning, 10-040-CM) supplemented with 10% FBS and 1% P/S. NIH 3T3 cells were cultured in DMEM supplemented with 10% calf serum (CS) (HyClone,SH3008703) and 1% P/S. For experiments, cells were seeded in 96-well or 384-well plates coated with fibronectin (MilliporeSigma, FC01010MG) diluted to 10 µg/mL in PBS. HeLa cells were seeded at a density of 7.5 x 10^3^ (96-well) or at a density of 4 x 10^3^ (384-well). HEK 293T and Beas2B cells were seeded at a density of 25 x 10^3^ (96-well) and 2.5 x 10^3^ (96-well) cells per well, respectively.

### Reagents and inhibitors

EML4-ALK inhibition was achieved using crizotinib (1 µM, Sigma-Aldrich, PZ0191) diluted in DMEM lacking FBS.

## Plasmid design and assembly

Cloning was performed using either PCR and scarless DNA assembly using NEBuilder® HiFi DNA Assembly Master Mix (New England Biolabs (NEB), #E2621); blunt end ligation using NEB T4 ligase (NEB, #M0202) or NEB KLD (NEB, #M0554S); or Golden Gate cloning using BbsI (NEB, R3539S). EML4-ALK variants were cloned downstream of (C-terminal to) GFP or mCh under control of a CMV promoter. Similarly, synthetic RTK multimers were cloned downstream of GFP or mCh under control of a CMV promoter. Grb2-GFP, CluMPS and the panel of GFP-tagged RTK fusions were described previously^4,12,13^. For endogenous tagging of GRB2 with GFP, donor DNA plasmid was constructed by integrating GFP in between two 500 nucleotide homology arms that flanked the stop codon of GRB2 into a pBlueScript backbone (pBS-HA(GRB2)-GFP) using HiFi DNA Assembly Master Mix. Donor DNA was prepared for transfection into HeLa cells by digesting the plasmid with NotI-HF (NEB, #R3189S) and EcoRI-HF (NEB,#R3101S) which were encoded in the 5’ and 3’ of the donor DNA sequence. Following digestion donor DNA was gel purified using Zymoclean Gel Recovery Kit (Zymo, #D4002). A previously reported 20 nucleotide single guided RNA sequence (CTTAGACGTTCCGGTTCACG)^6^ was integrated into the pX330 spCas9-mSA (Addgene# 113096) using Golden Gate cloning. For integration of Erk-KTR-iRFP into HeLa Grb2:GFP cells using Super PiggyBac Transposase, Erk-KTR-iRFP was integrated into pBRPB Cag-mCherry-IP (Addgene, plasmid # 106333) by excising mCherry out of the plasmid using PCR and integrating Erk-KTR-iRFP using HiFi Assembly Master Mix. For stable expression of synthetic ALK constructs and EML4-ALK(V1) in 3T3 cells GFP-1mer-ALK, GFP-2mer-ALK, GFP-2mer-ALK(KD) and GFP-EML4-ALK(V1) were integrated into a pHR lentivarial backbone using HiFi Assembly Master Mix.

### Transient transfection and genome integration

Exogenous expression of RTK fusion constructs was done using FuGENE 4K Transfection Reagent (Promega, E5912). In short, 25-100 ng of plasmid were mixed with 0.3-0.6 µL of FuGENE 4K transfection reagent in 10 µL of OptiMEM (Gibco, 31985-070). After 12 min incubation at room temp, the transfection mix was added to culture media lacking P/S. For transfections in a 96-well plate, 10 µL of transfection mix were added to 200 µL of media per well. For transfections in 384-well plates, 2.5 µL of transfection mix were added to 50 µL media per well. Four to six hours after transfection, cells were washed with starvation media (DMEM or RPMI with P/S but without FBS). For endogenous tagging of GRB2 with GFP, HeLa cells were co-transfected with donor DNA and pX330 spCas9-mSA-sgRNAGRB2 using FuGene 4K Transfection Reagent. Proper tagging was validated through Western blot. Stable expression of EKR-KTR-iRFP in HeLa GRB2:GFP cells was achieved by co-transfecting Super PiggyBac Transposase (Systems Biosciences, #PB210PA-1) using Fugene 4K.

### Lentiviral packaging and cell line generation

Stable cell lines were made using lentivirus. Constructs were packaged into lentiviral particles by cotransfecting pHR transfer plasmids with pCMV-dR8.91 (Addgene, Catalog#12263) and pMD2.G (Addgene, Catalog#12259) into HEK293T cells using the calcium phosphate transfection method previously described^35^. Following 2 days of media collection, media containing virus was centrifuged at 200 x g for 2 min and supernatant was filtered using a 0.45-µm filter. 500 µL of each virus was then added to 1 x 10^5^ cells in one well of a 6-well plate.

### Immunostaining

Cells were treated for 10 min with 4% paraformaldehyde (PFA) (Electron Microscopy Sciences, 15710) for chemical fixation. Cells were then permeabilized using 0.5% Triton X-100 in PBS for 10 min. When probing for Erk phosphorylation, cells were subsequently incubated in 100% methanol at-20°C for 10 min. Cells were then blocked using 1% Bovine Serum Albumin (BSA) (Fisher, BP9706100) diluted in Phosphate Buffer Saline (PBS) for 1 h at room temperature. Cells were then incubated with primary antibody overnight at 4°C. Primary antibodies: phospho-p44/42 MAPK (ERK1/2) (Thr202/Tyr204) diluted 1:400, Cell Signaling #4370; phospho-ALK receptor (Tyr 1507) diluted 1:200, Cell Signaling #14678; phospho-RET receptor (Tyr 1096) diluted 1:200, Invitrogen # PA5-105796. Cells were then washed 5X with 0.1% Tween-20 (Fisher, 9005-64-5) in PBS (PBS-T). Cells were incubated in secondary antibody 1:500 and with 4,6-diamidino-2-phenylindole (DAPI; ThermoFisher Scientific #D1306, 300 nM) for 1 hr at room temperature and washed as previously 5X with 0.1% PBS-T. Secondary antibodies: antibody IgG (H+L) Cross-Adsorbed Goat anti-Rabbit, DyLight™ 488, Invitrogen #35553; Goat anti-Rabbit IgG (H+L) Cross-Adsorbed Secondary Antibody, DyLight™ 650, Invitrogen #SA510034.

### Fixed-cell imaging

Following immunostaining, cells were imaged with a 40X air or 60X oil objective using a Nikon Ti2-E microscope equipped with a Yokagawa CSU-W1 spinning disk, 405/488/561/640 nm laser lines, an sCMOS camera (Photometrics), and a motorized stage.

### Live-cell imaging

Live-cell imaging was performed using a Nikon Ti2-E microscope equipped with a Yokagawa CSU-W1 spinning disk, 405/488/561/640 nm laser lines, an sCMOS camera (Photometrics), and a motorized stage equipped with an environmental chamber that maintained temperature at 37°C and 5% CO_2_. For experiments measuring Grb2 localization over time, HeLa Grb2:GFP cells expressing Erk-KTR-iRFP were transiently transfected with mCh-EML4-ALK(V1). 24 hrs after transfection, cells were incubated with 5 µg/mL Hoechst 33342 Hydrochloride (Cayman 15547) for 15 min and were then washed with serum-free DMEM. Images were acquired each minute for 30 min, starting from the moment of ALKi treatment, using a 40X air objective. GRB2 localization in cells expressing synthetic ALK constructs was imaged following 24 hrs of transfection using a 40X air objective.

### Condensate and diffuse RTK fusion quantification

Cell body and cell nucleus segmentation was performed using Cellpose^36^, a generalist algorithm for cellular segmentation. RTK fusion condensates were segmented using a custom MATLAB script described previously^13^. Cell body, nucleus, and RTK fusion condensate masks were then processed along with raw images to obtain intensity and size metrics using CellProfiler^37^. For pALK compartmentalization analysis, pALK intensity readings were background-subtracted using pALK staining intensity from untransfected HeLa cells. For Erk-KTR analysis, the cytoplasmic intensity was obtained by measuring a 5 pixel ring surrounding the cell’s nucleus. Cell expression was gated into bins using half logs unless otherwise specified in the legend. Data was then exported to RStudio for data wrangling and visualization using the tidyR packages^38^.

### Soft agar colony formation assay

To assess transformation capacity of the synthetic ALK constructs, 5 x 10^3^ cells stably expressing GFP-1mer-ALK, GFP-2mer-ALK, GFP-2mer-ALK(KD), or GFP-EML4-ALK(V1) were seeded in 1mL of DMEM supplemented with 10% CS, 1%P/S and 0.3% agarose on top of a 2 mL layer of DMEM supplemented with 10% CS, 1%P/S and 0.6% agarose in a 6 well plate that had been allowed to set while preparing the cells for seeding. Each well then received 1 mL of DMEM supplemented with 10% CS and 1% P/S. Media was exchanged every 7 days. After 4 weeks, cells were fixed with 4% PFA and stained with 0.05% crystal violet in PBS for 1 hr. Colonies were imaged using a 4X objective using a Nikon Ti2E epifluorescence microscope.

### 3T3 xenograft assay

Xenografts were performed by suspending 1M 3T3 cells stably expressing GFP-EML4-ALK, 2mer-ALK, 1mer-ALK(CA), or 2mer-ALK(KD) in 400µl of PBS and mixed with 400µl of VitroGel Hydrogel Matrix (The Well Bioscience; VHM01). 200µl of cell suspension (2.5 x 10^5^ cells) were then injected subcutaneously into the flank of three NSG mice per construct. Tumor size was calculated using 𝑉 = (𝑙𝑒𝑛𝑔𝑡ℎ × 𝑤𝑖𝑑𝑡ℎ^!^)/2.

## Fluorescence correlation spectroscopy

For in-cell fluorescence correlation spectroscopy (FCS) measurements, HEK 293T cells were plated in fibronectin coated 8-well NUNC chambers (Thermo Scientific, Rochester, NY) and transfected with GFP or mutant GFP-RTK constructs. Cell media was exchanged with phenol red-free FluoroBrite media (Thermo Scientific, cat. A189701) approximately 30 minutes prior to imaging and FCS measurements. A portable stage-top incubator was used during measurements to maintain the environment at 37 °C and 5% CO_2_.

Fluorescence lifetime imaging (FLIM) and FCS measurements were carried out on a PicoQuant MicroTime 200 fluorescence microscope system equipped with a FLIMbee galvo scanner (PicoQuant, Berlin, Germany) and Nano-ZL 100 piezo scanning stage (Mad City Labs, Madison, WI) to allow for imaging in three dimensions as described previously^13^. Briefly, a 482 nm, 40 MHz laser was set to 65% of its maximum output and focused by a 60X Plan-Apo/1.4-NA water-immersion objective (Olympus, Tokyo, Japan). Fluorescence emission was collected through the objective and focused onto a 150 μm diameter aperture, then directed through a 525 ± 35 nm band-pass filter and collected by an avalanche photodiode detector. Image acquisition and data collection were done using PicoQuant’s accompanying software SymPhoTime 64.

To select for cells with appropriate concentrations of fluorescent protein for FCS measurements, a large field (261 μm x 261 μm) of cells were imaged with 50 μs per pixel dwell time and laser power as noted above. The excitation volume was then moved to the interior of an individual cell and the count rate measured; cells with count rates within the range of 60–500 kHz were selected to move forward. For a selected cell, an image in the X/Y plane was first acquired, followed by an image in the X/Z or Y/Z plane. The X/Z or Y/Z image was used to determine the Z position of the excitation volume in the cell cytoplasm, allowing us to avoid measuring too close to the plasma membrane or nucleus. Generally, 3 to 5 FCS measurements were acquired at different X/Y locations inside each cell. For each FCS measurement, ten autocorrelation curves of ten seconds each were collected and averaged together. Using a lab-written script in MATLAB, the averaged autocorrelation curves were fit to a diffusion model for a single fluorescent species undergoing subdiffusive motion in a three-dimensional Gaussian volume with molecular crowding, with fits weighted by the inverse square of the standard deviation. The autocorrelation function for this anomalous diffusional model is described by Equation 1.

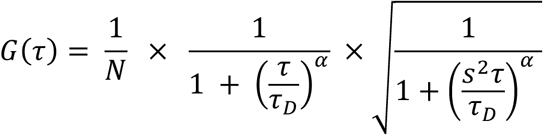

where *N* is the average number of molecules in the focal volume, *s* is the ratio of the radial to axial dimensions of the focal volume, is the translational diffusion time, and a reflects the degree of molecular crowding in the cellular environment^39–41^. For each FCS measurement, the molecular brightness of the fluorescent species – expressed as counts per molecule (CPM) – was calculated by dividing the average intensity in kHz by *N*.

## Supplemental Figures

**Supplementary Figure 1.**
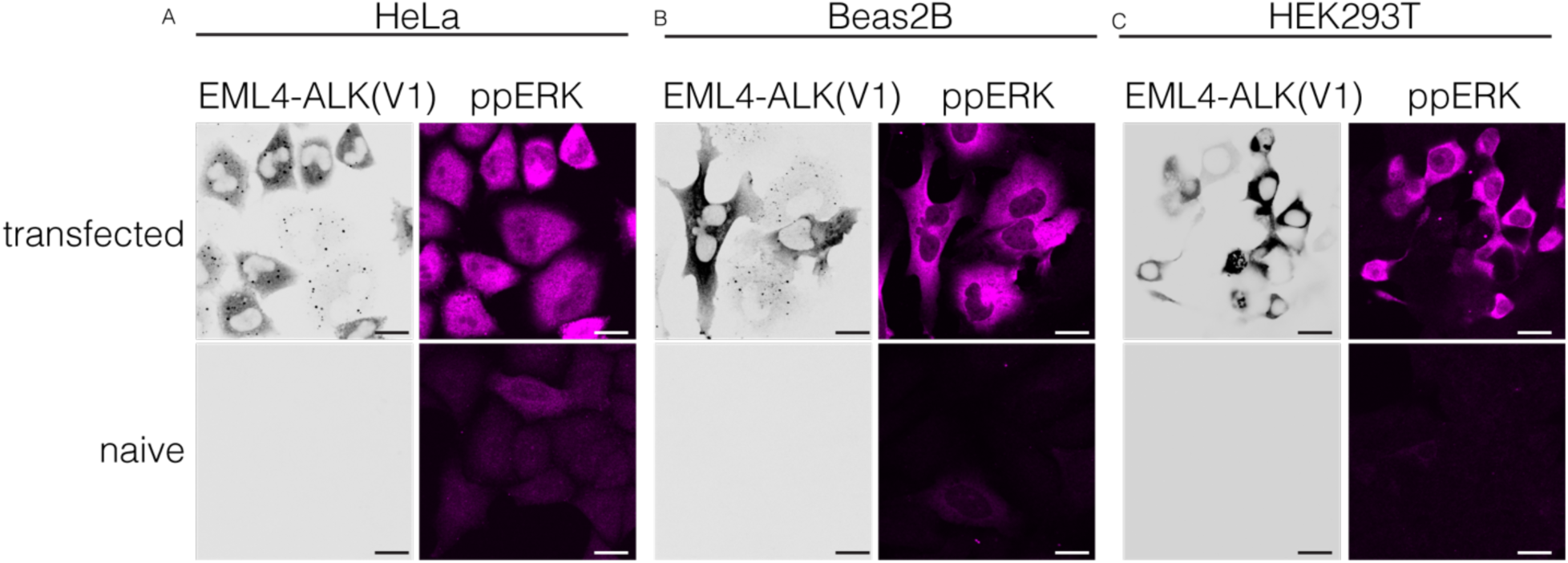
EML4-ALK expression activates Erk signaling in HeLa, Beas2B, and HEK 293T cells despite differences in condensation. Representative images of pERK immunostaining in HeLa, Beas2B, and HEK 293T cells expressing EML4-ALK (V1). Top row: transfected with mCh-EML4-ALK(V1). Bottom row: untransfected. Scale bars = 20 µm.

**Supplementary Figure 2.**
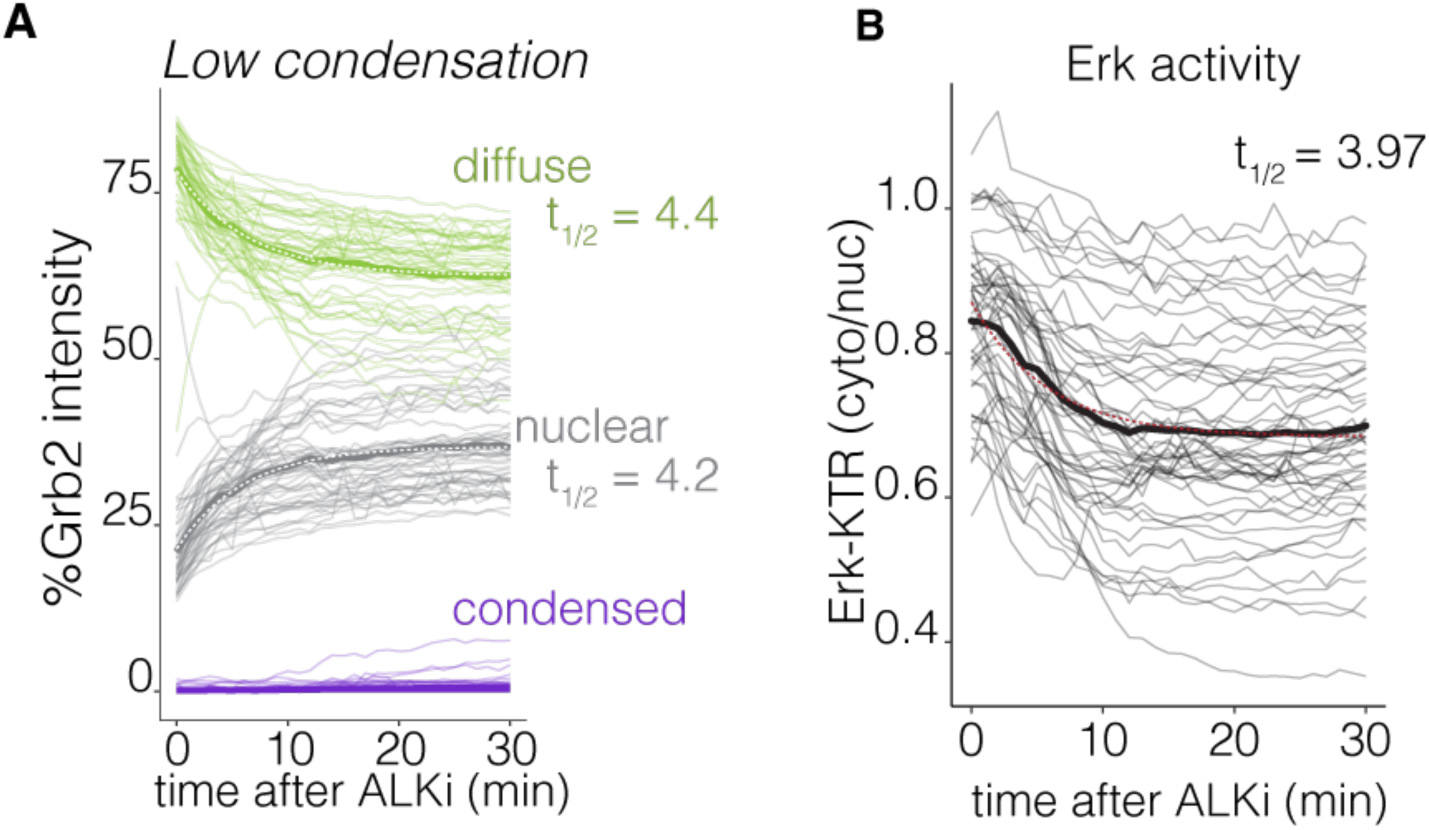
Comparison of GRB2 localization and Erk activity after ALK inhibition in cells lacking EML4-ALK condensates. A) Tracking GRB2:GFP fluorescence intensity in pixels corresponding to diffuse or condensed EML4-ALK(V1), or in the nucleus, following ALKi (crizotinib, 1 µM) in HeLa:GRB2-GFP cells expressing exogenous EML4-ALK(V1). Traces correspond to cells with low condensation (<0.2% area in condensates). Data of high condensation cells are shown in **Figure 3L**. B) ERK-KTR activity (cytoplasmic to nuclear intensity ratio) following ALKi treatment of cells in (A). Light traces represent single cells. Dark traces represent the mean of single cell traces. Dashed lines represent the best fit single exponential decay, which was used to calculate t_1/2_.

**Supplementary Figure 3.**
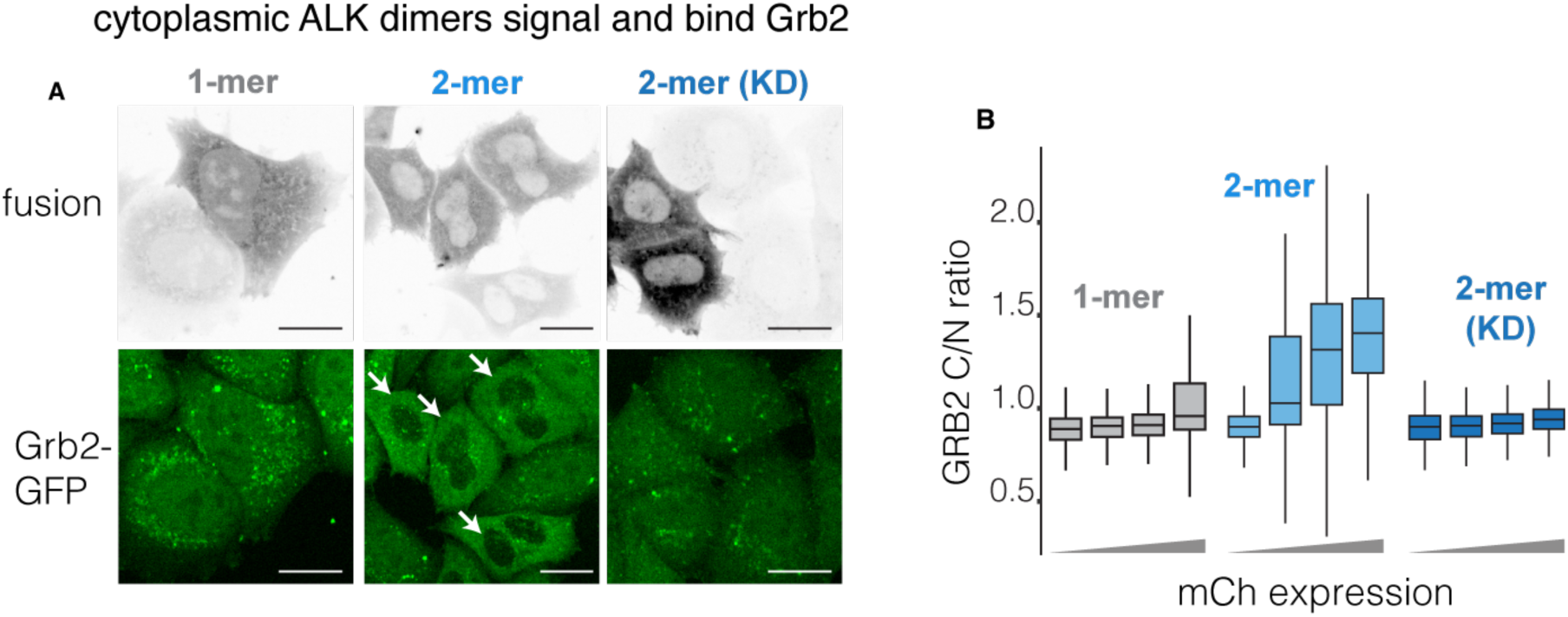
Active 2mer-ALK recruits GRB2 in the cytoplasm. A) Representative images of the indicated constructs expressed in HeLa GRB2:GFP cells. KD = kinase dead. White arrows highlight the GRB2:GFP distribution and nuclear exclusion in cells that were transfected with the signaling-competent 2-mer. Scale bars = 20 µm. B) GRB2:GFP cyto/nuc ratio of 1-mer, 2-mer, 2-mer(KD) expressed in HeLa cells, as a function of expression. n = 398-492 (1-mer), 366-432 (2-mer), 663-813(2-mer (KD)) cells.

**Supplementary Figure 4.**
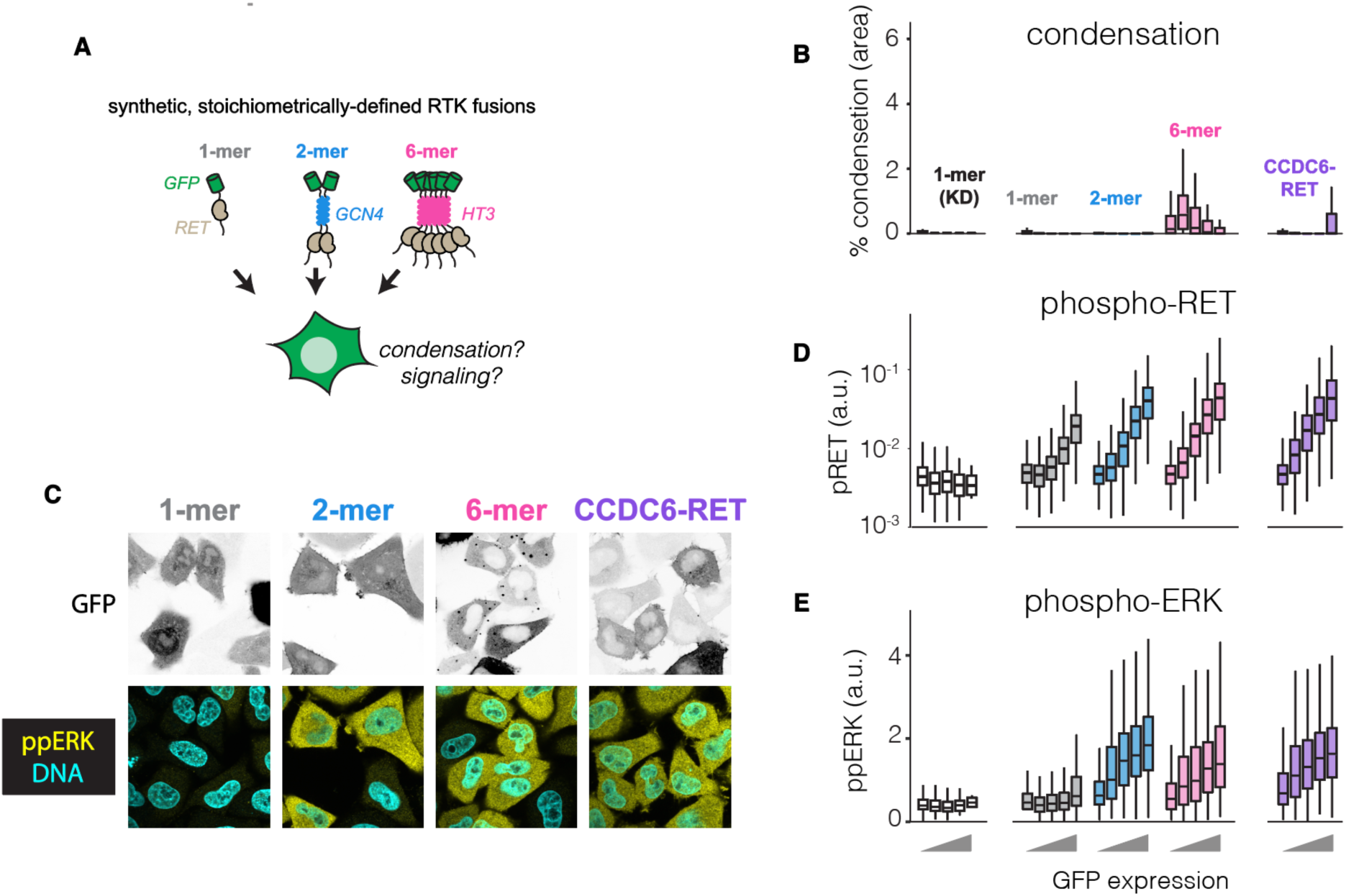
RET fragment dimerization is sufficient for activation and RAS/ERK signaling. A) Synthetic, stoichiometrically defined multimers of the RET intracellular domain were made to assess the contribution of multivalency to activation of the RET fusions and downstream ERK signaling. B) Condensation of RET multimers and CCDC6-RET. RET 6mer forms condensates over a range of expression, while RET 1mer and 2mer do not. A control kinase-dead 1mer also does not form condensates. n = ∼50-1000 cells per construct per expression bin. C) Representative images of RET 1mer, 2mer, 6mer and CCDC6-RET. 1mer-RET does not activate ERK, while 2mer-, 6mer-, and CCDC6-RET result in ERK phosphorylation. Scale bars = 20 µm. D) Quantitation of phosphorylation of the RET tail as a function of fusion concentration. n = ∼200-2000 cells per construct per expression bin. E) Quantitation of ERK phosphorylation as a function of RET fusion concentration. n = ∼200-2000 cells per construct per expression bin. As for ALK fusions (**Figure 4**), RET fusion dimers are sufficient to trigger RET phosphorylation and ERK signaling in the absence of condensation.

**Supplementary Figure 5.**
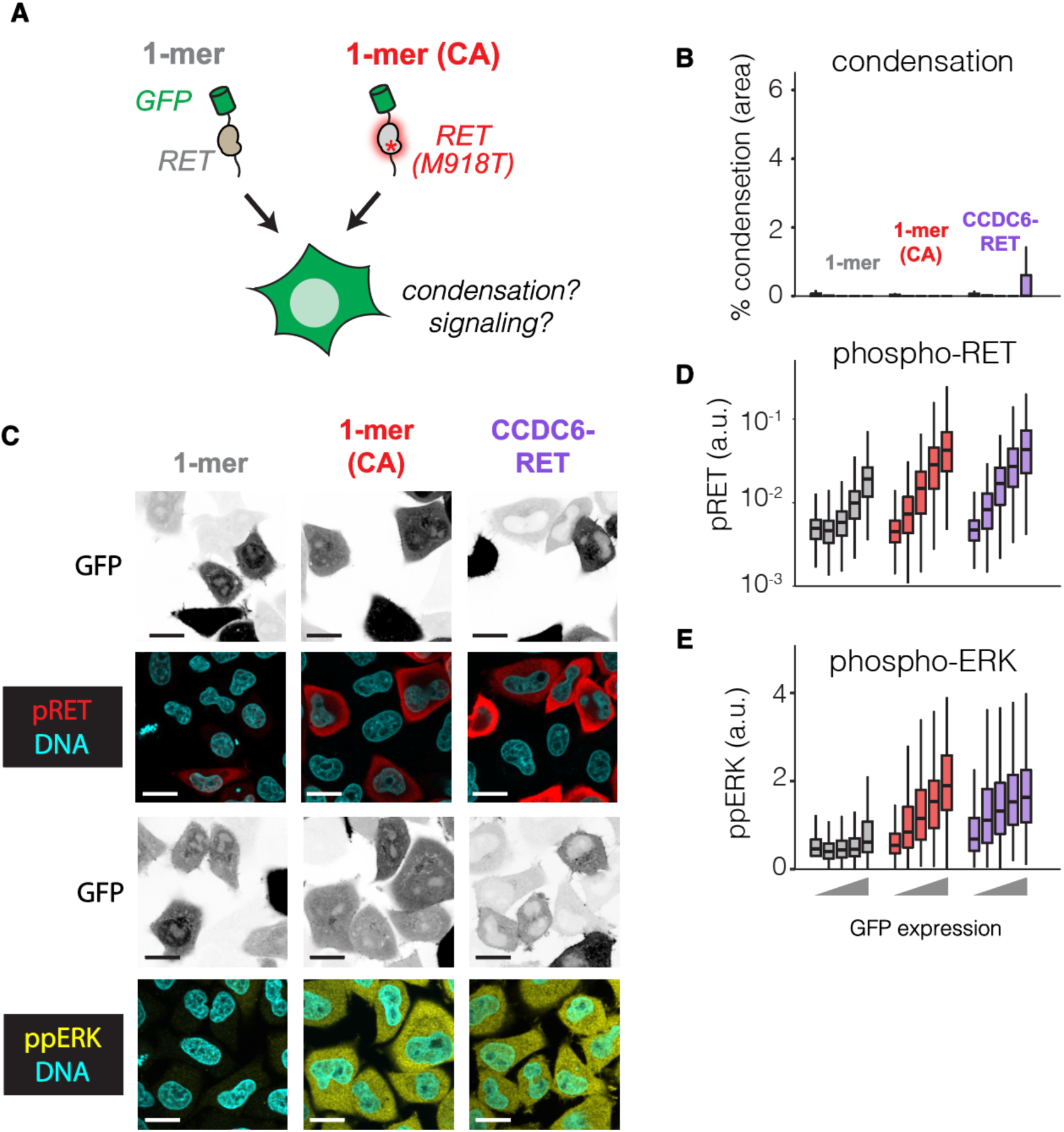
Constitutively active RET monomers achieve kinase activation and trigger RAS-ERK signaling. A) Monomers of the wildtype or constitutively active (CA) RET fragments were made to assess if RET phosphorylation could activate ERK signaling in the absence of multimers and/or condensates. B) Quantification of condensation of 1mer-RET, 1mer-RET(CA), and fusion oncoprotein CCDC6-RET. Neither RET 1mer forms condensates at any expression level. n = ∼50-1000 cells per construct per expression bin. C) Representative images of RET constructs and corresponding phospho-RET or pERK immunofluorescence. 1mer-RET(CA) triggers pRET and pERK comparably to CCDC6-RET, whereas 1mer-RET does not. Scale bar = 20 µm. D) Quantification of pRET as a function of fusion expression. n = ∼200-2000 cells per construct per expression bin. E) Quantification of pERK signaling as a function of RET fusion expression. n = ∼200-2000 cells per construct per expression bin.

## Supplementary Movie Captions

**Supplementary Movie 1. GRB2 localization and Erk activity after ALKi treatment.** HeLa:Grb2-GFP cells co-expressing mCh-EML4-ALK(V1) and Erk-KTR-iRFP imaged after treatment with ALKi (crizotinib, 1 µM). Time in hh:mm.

